# Structural basis for CFTR inhibition by CFTR_inh_-172

**DOI:** 10.1101/2023.10.11.561899

**Authors:** Paul Young, Jesper Levring, Karol Fiedorczuk, Scott C. Blanchard, Jue Chen

## Abstract

The cystic fibrosis transmembrane conductance regulator (CFTR) is an anion channel that regulates electrolyte and fluid balance in epithelial tissues. Whereas activation of CFTR is vital to treating cystic fibrosis, selective inhibition of CFTR is a potential therapeutic strategy for secretory diarrhea and autosomal dominant polycystic kidney disease (ADPKD). Although several CFTR inhibitors have been developed by high-throughput screening, their modes of action remain elusive. In this study, we determined the structure of CFTR in complex with the inhibitor CFTR_inh_- 172 to 2.7 Å resolution by cryogenic electron microscopy (cryo-EM). We observe that CFTR_inh_- 172 binds inside the pore near transmembrane helix 8 (TM8), a critical structural element that links ATP hydrolysis with channel gating. Binding of CFTR_inh_-172 stabilizes a conformation in which the chloride selectivity filter is collapsed and the pore is blocked from the extracellular side of the membrane. Single molecule fluorescence resonance energy transfer (smFRET) experiments indicate that CFTR_inh_-172 inhibits channel gating without compromising nucleotide-binding domain (NBD) dimerization. Together, these data show that CFTR_inh_-172 acts as both a pore blocker and a gating modulator, setting it apart from typical ion channel inhibitors. The dual functionality of CFTR_inh_-172 reconciles previous biophysical observations and provides a molecular basis for its activity.

**Significance statement:** The pathogenesis of secretory diarrhea and autosomal dominant polycystic kidney disease involves hyperactivation of the CFTR ion channel. CFTR inhibitors, including the small-molecule CFTR_inh_-172, have been developed as therapeutic candidates to treat these diseases. This study offers a structural understanding of CFTR_inh_-172’s mode of action, clarifying its dual inhibitory role as both a pore blocker and gating modulator. The molecular description of how CFTR_inh_-172 interacts with CFTR provides a structural foundation to its specificity and efficacy. Furthermore, the observation that CFTR inhibitors and potentiators both interact with TM8 strengthens the notion that this helix serves as an allosteric link between the ATPase site and the channel gate, and is therefore a hotspot for pharmacological modulation.

## INTRODUCTION

The cystic fibrosis transmembrane conductance regulator (CFTR) is an anion channel expressed in the apical membrane of epithelial cells (1, 2). Loss-of-function mutations in the *cftr* gene cause widespread salt and fluid dysregulation that leads to the autosomal recessive disease cystic fibrosis (1). By contrast, hyperactivation of CFTR is central to pathogenesis in secretory diarrhea and autosomal dominant polycystic kidney disease (ADPKD) (3–10). In both secretory diarrhea and ADPKD, cyclic AMP (cAMP) accumulation activates protein kinase A (PKA) (11–15). PKA-phosphorylation in turn activates CFTR (11, 12, 16–18), leading to excess fluid accumulation in the intestinal lumen and renal cysts, for secretory diarrhea and ADPKD, respectively (19, 20). CFTR hyperactivation has also been implicated in the pathogenesis of non-alcoholic steatohepatitis (NASH) (21).

Despite functioning as an ion channel, CFTR belongs to the superfamily of ATP-binding cassette (ABC) transporters. It is comprised of two transmembrane domains (TMDs) and two nucleotide-binding domains (NBDs) that are common to all ABC transporters, along with a cytosolic regulatory (R) domain specific to CFTR (22, 23). CFTR’s activity is regulated at two levels. Phosphorylation by PKA releases the auto-inhibition imposed by the unphosphorylated R domain (24, 25). Once phosphorylated, ATP-binding promotes NBD dimerization and pore opening, whereas ATP hydrolysis leads to pore closure (26).

Substantial effort has been devoted to understanding CFTR in the context of cystic fibrosis. Disease-causing mutations have been extensively characterized, and small molecule drugs that potentiate gating or correct folding of CFTR have been successfully developed for clinical use (27–38). By contrast, relatively few studies have addressed CFTR hyperactivation. Although secretory diarrhea is the second leading cause of death in children under 5 worldwide (20, 39, 40), and ADPKD is the most common inherited cause of end-stage renal disease (ESRD) (19), both conditions lack broadly effective, generalizable pharmacological treatments. Despite the therapeutic potential of CFTR inhibitors and although several small-molecule CFTR inhibitors have been identified (5, 41–43), their mechanisms and sites of action remain poorly understood.

In this study, we investigate the mechanism of CFTR_inh_-172, a highly specific and efficacious CFTR inhibitor developed in the Verkman laboratory (5). This inhibitor was shown to block cholera toxin-induced intestinal fluid secretion (5) and suppress cyst growth in animal models of polycystic kidney disease (3). Substitution of pore-lining residues reduced the potency of CFTR_inh_-172, suggesting that it directly binds to CFTR (44). However, several studies have also suggested that CFTR_inh_-172 acts as a gating modulator rather than a classical pore-blocker (45, 46). Using cryo-EM, smFRET, and electrophysiology, we have discovered that CFTR_inh_-172 binds within the pore, stabilizing the transmembrane helices in a nonconductive conformation without obstructing NBD dimerization. These findings enable us to propose a mechanism for CFTR_inh_-172 that reconciles previous conflicting observations.

## RESULTS

### CFTR_inh_-172 inhibits wild-type CFTR and the “locked-open” CFTR (E1371Q)

First, we characterized the effects of CFTR_inh_-172 in excised inside-out membrane patches (Figure 1A-B). Consistent with previous work (46), application of 10 µM CFTR_inh_-172 reduced the macroscopic current of wild-type (WT) CFTR by 96%. A comparable level of inhibition was observed with the hydrolysis-deficient CFTR (E1371Q), which has an open dwell time 1000-fold longer than that of WT CFTR (47). Next purified CFTR was reconstituted into a planar lipid bilayer and activated by PKA. Single channel currents for WT CFTR and CFTR (E1371Q) were measured in the presence of 3 mM ATP with or without 10 µM CFTR_inh_-172 (Figure 1C). CFTR_inh_-172 reduced the open probability of WT CFTR from 0.21 ± 0.05 (mean and standard error) to 0.007 ± 0.003, whereas that of CFTR (E1371Q) decreased from 0.79 ± 0.03 to 0.0011 ± 0.0004 (Figure 1D). In agreement with a previous observation that CFTR_inh_-172 affects both the open dwell time and closed dwell time (46), we observed a large effect on closed dwell time and a fivefold reduction in mean open dwell time, from 487 ± 92 ms to 109 ± 14 ms for WT CFTR (Figure 1E). The similar responses of WT CFTR and CFTR (E1371Q) to CFTR_inh_-172 suggest a comparable mechanism of inhibition and establish CFTR (E1371Q) as a suitable system for structural studies with this inhibitor.

**Figure 1.**
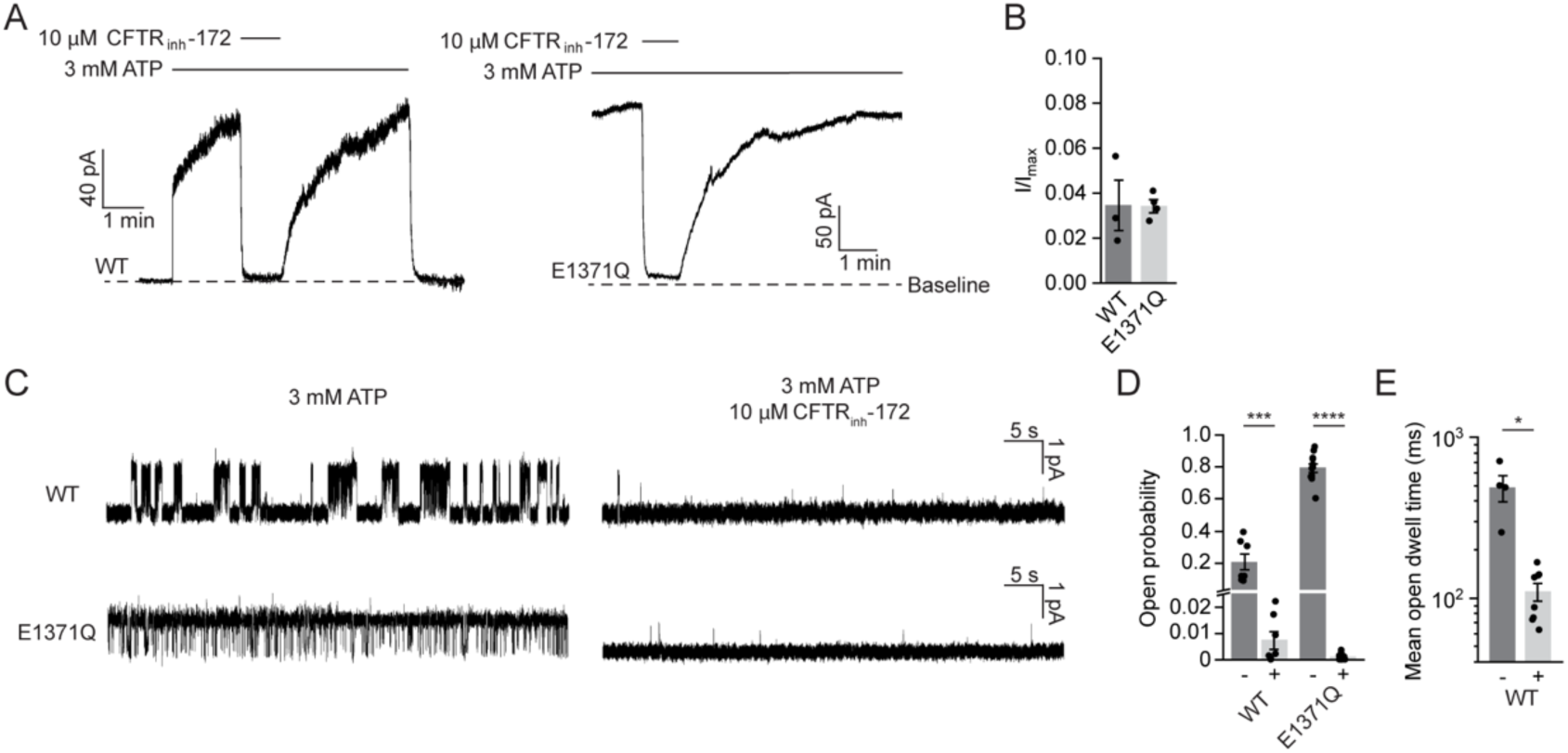
CFTR_inh_-172 inhibits WT and E1371Q variant CFTR. (**A**) Example macroscopic current traces showing inhibition of WT CFTR and CFTR (E1371Q) by CFTR_inh_-172 in inside-out excised patches. CFTR was fully phosphorylated by PKA in the presence of 3 mM ATP before the displayed recordings. (**B**) Currents with 10 µM CFTR_inh_-172 relative to currents without CFTR_inh_-172.. Data represent means and standard errors of 3 (WT) or 4 (E1371Q) patches. Individual data points are displayed as dots. (**C**) Example single-channel recordings of PKA-phosphorylated WT and CFTR (E1371Q) reconstituted in synthetic lipid bilayers with and without CFTR_inh_-172. Recordings were made at -150 mV. Upward deflections correspond to opening. (**D**) Open probabilities of WT CFTR and CFTR (E1371Q) with and without CFTR_inh_-172. Data represent means and standard errors of 7 (WT, with or without CFTR_inh_-172), 11 (E1371Q, without CFTR_inh_-172), or 9 (E1371Q, with CFTR_inh_-172) bilayers. Statistical significance was tested by one-way analysis of variance (\*\*\**p* = 2×10^-4^, \*\*\*\**p* < 10^-10^). (**E**) Mean open dwell time of WT CFTR with and without CFTR_inh_-172. Data represent means and standard errors of 4 (without CFTR_inh_-172), or 8 (with CFTR_inh_-172) bilayers. Statistical significance was tested using two-tailed Student’s t-test (\**p* = 0.025).

### CFTR_inh_-172 binds within the pore

To identify the binding site of CFTR_inh_-172, we determined the cryo-EM structure of inhibitor-bound CFTR using phosphorylated CFTR (E1371Q) (Figure 2, Figure S1). The structure was determined to an overall resolution of 2.7 Å. In the presence of CFTR_inh_-172 and ATP, CFTR (E1371Q) adopts an NBD-dimerized, pore-closed conformation distinct from any previously observed structures (Figure 2A). The density for CFTR_inh_-172, as strong as the protein main chain atoms, was observed inside the pore, at a position corresponding to the membrane outer leaflet (Figure 2A). The chemical structure of CFTR_inh_-172 can be divided into three rings (Figure 2B): a central thiazolidine ring (Ring B) with a (3-trifluoromethyl)phenyl substitution at position 3 (Ring A) and a (4-carboxyphenyl)methylene substitution at position 5 (Ring C). The density shows well-defined features corresponding to the trifluoromethyl phenyl and the heavy sulfur atoms in the thiazolidine ring (Figure 2B). The density for Ring C is not as well-defined, indicating that this moiety may be mobile.

**Figure 2.**
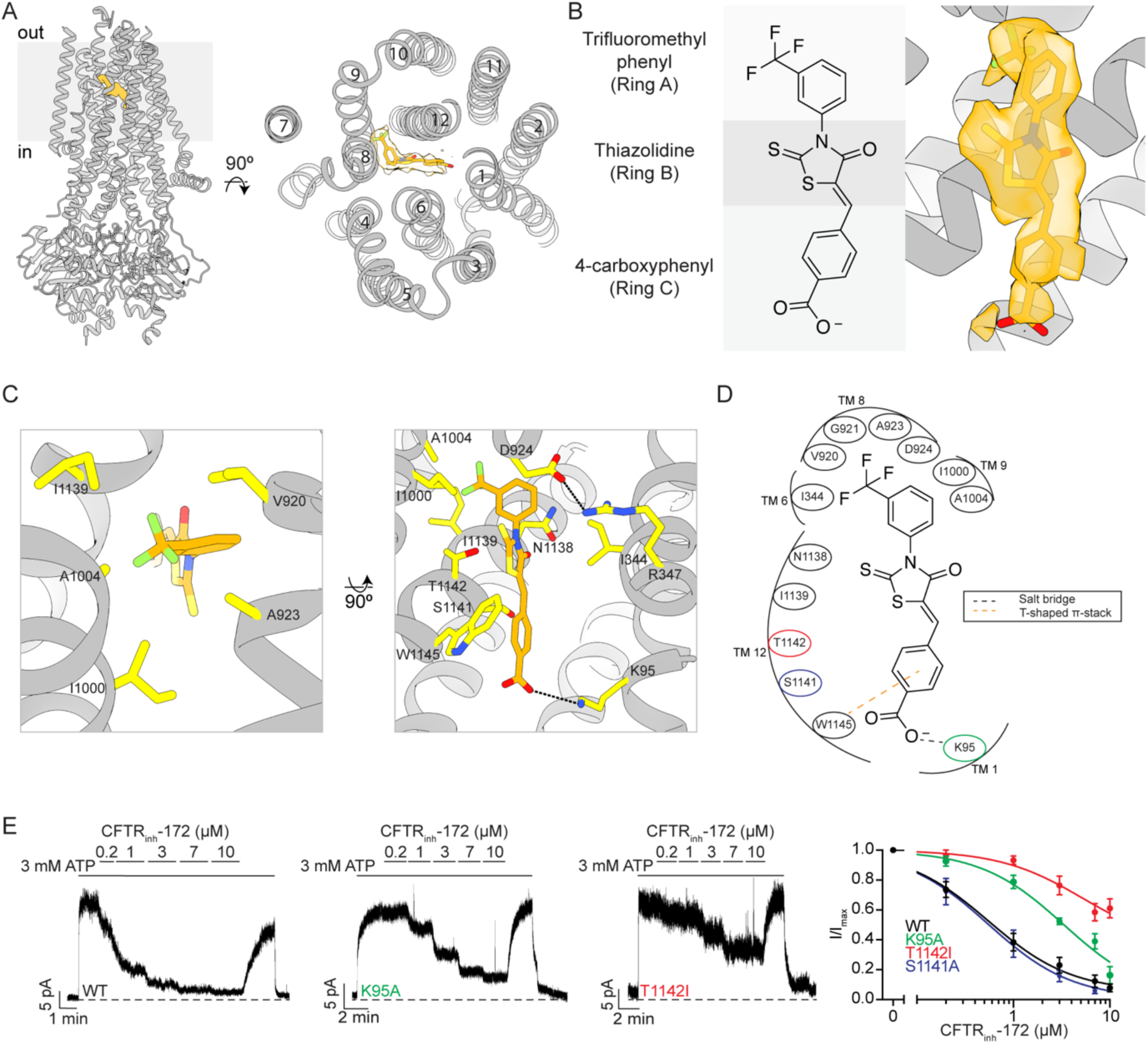
CFTR_inh_-172 makes specific interactions with the CFTR inner vestibule. (**A**) (left) Structure of phosphorylated ATP-bound CFTR (E1371Q) bound to CFTR_inh_-172. CFTR_inh_-172 is shown as an orange stick model surrounded by cryo-EM density. (right) A view of the structure looking down the long axis of the CFTR pore with CFTR_inh_-172 modeled as orange sticks surrounded by cryo-EM density. (**B**) (left) Structure of CFTR_inh_-172 with the three functional rings delineated by gray rectangles. (right) Structure of CFTR_inh_-172 modeled into its binding site within the inner vestibule surrounded by cryo-EM density. (**C**) Interacting residues within 4.5 Å of CFTR_inh_-172. (left) The hydrophobic pocket of the CFTR_inh_-172 binding site. (right) The solvent-exposed pocket of the CFTR_inh_-172 binding site. CFTR_inh_-172/K95 and R347/D924 salt bridges are shown as black dashed lines. (**D**) Schematic drawing of the CFTR-inhibitor interactions. All residues within 4.5 Å of the inhibitor are depicted. Residues substituted in inside-out path-clamp electrophysiology are indicated with colored circles. (**E**) Example macroscopic current traces showing titration of CFTR_inh_-172 onto WT, K95A, or T1142I CFTR in inside-out excised patches. CFTR was fully phosphorylated by PKA in the presence of 3 mM ATP before CFTR_inh_-172 titration. (right) Dose-response curves for CFTR_inh_-172 binding site variants. The mean current in the presence of 3 mM ATP alone was used to normalize current elicited at each CFTR_inh_-172 concentration. Dose-response curves were fitted with the Hill equation to estimate *IC_50_* values for each variant. Hill coefficients were fixed to 1. Each data point represents mean and standard error determined from three to five patches.

CFTR_inh_-172 binds within the CFTR pore and interacts with both TMDs (Figure 2A). The three rings of the compound form an elongated shape wedged between TMs 1, 6, 8, 9, and 12. Whereas Ring A and Ring B are completely buried, Ring C is exposed to the aqueous ion-conduction pathway (Figure 2A). With this structure, we can now fully understand previous structure-activity-relationship (SAR) analyses (5, 48). Specifically, these studies showed that the trifluoromethyl group (CF_3_) is essential for inhibition and that the addition of polar substituents or removal of CF_3_ on Ring A diminished activity (48). The highest potency is achieved when CF_3_ is at position 3 of Ring A, as in CFTR_inh_-172 (5, 48). In the structure, the trifluoromethyl group fits snugly into a hydrophobic cavity, establishing van der Waals interactions with five hydrophobic residues on TMs 6, 8, 9, and 12 (Figure 2C-D). If the CF_3_ substitution were to occur at positions 2 or 4, many of these interactions would be lost, resulting in lower inhibitory potency, as observed (48). The SAR studies further demonstrate the significance of negative or polar substitutions on Ring C (48). Indeed, we observe that this ring is positioned within a spacious, solvent-exposed cavity surrounded by numerous charged and polar residues, including K95 on TM1, Q353 on TM6, as well as N1138, S1141, and T1142 on TM12 (Figure 2C-D). The carboxy group on CFTR_inh_-172 forms a salt bridge with K95 (Figure 2C-D). Consistent with this observation, esterification or amidation of the carboxy group in CFTR_inh_-172 resulted in inactive compounds (48), presumably due to the loss of this interaction. The reciprocal change on CFTR, substitution of K95 with alanine, resulted in a nearly sevenfold decrease in potency, increasing the half maximal inhibitory concentration (*IC_50_*) of CFTR_inh_-172 from 0.6 ± 0.1 µM to 3.5 ± 0.9 µM (Figure 2E). Perturbation of the binding site by T1142I substitution increased the *IC_50_* to 5.3 ± 3.1 µM (Figure 2E). The effect of the T1142I substitution is most likely due to steric hindrance, as the side chain of T1142 is close to the methylene group at position 2 of Ring B (Figure 2C-D). The reciprocal modification on the inhibitor, adding a methyl at this position, was shown to increase the *IC_50_* to 8 µM (48). By contrast, alanine substitution of a non-interacting residue, S1141, had no effect on inhibitory potency (Figure 2E, Figure S2).

The structure also offers a molecular explanation for previous data showing that substituting R347 with alanine decreased the potency of CFTR_inh_-172 by over 30-fold, and that R347D substitution nearly eliminated its inhibitory effect (44). Although R347 does not interact with the inhibitor directly, it forms a salt bridge with D924, creating a surface against which the inhibitor is tightly packed (Figure 2C). Substitutions at position 347 are likely to modify the structure of the binding site, consequently reducing the inhibitory activity.

### CFTR_inh_-172 stabilizes a closed conformation of CFTR

Previous structural studies of human CFTR have revealed the conformational changes required for pore opening (Figure 3A). Dephosphorylated CFTR exhibits an NBD-separated conformation with an inner vestibule open to the cytosol but the pore closed off to the extracellular space (Figure 3A) (49). In the presence of ATP, phosphorylated CFTR (E1371Q) forms an NBD-dimerized conformation (50), in which the pore is open and a dehydrated chloride ion is bound at the selectivity filter near the extracellular entrance (see accompanying paper) (Figure 3A). A comparison of these two structures reveals that phosphorylation and ATP binding cause the NBDs and TMDs to move towards the central axis essentially as rigid bodies. However, local conformational changes of TMs 8 and 12 are also critical for CFTR gating (25).

**Figure 3.**
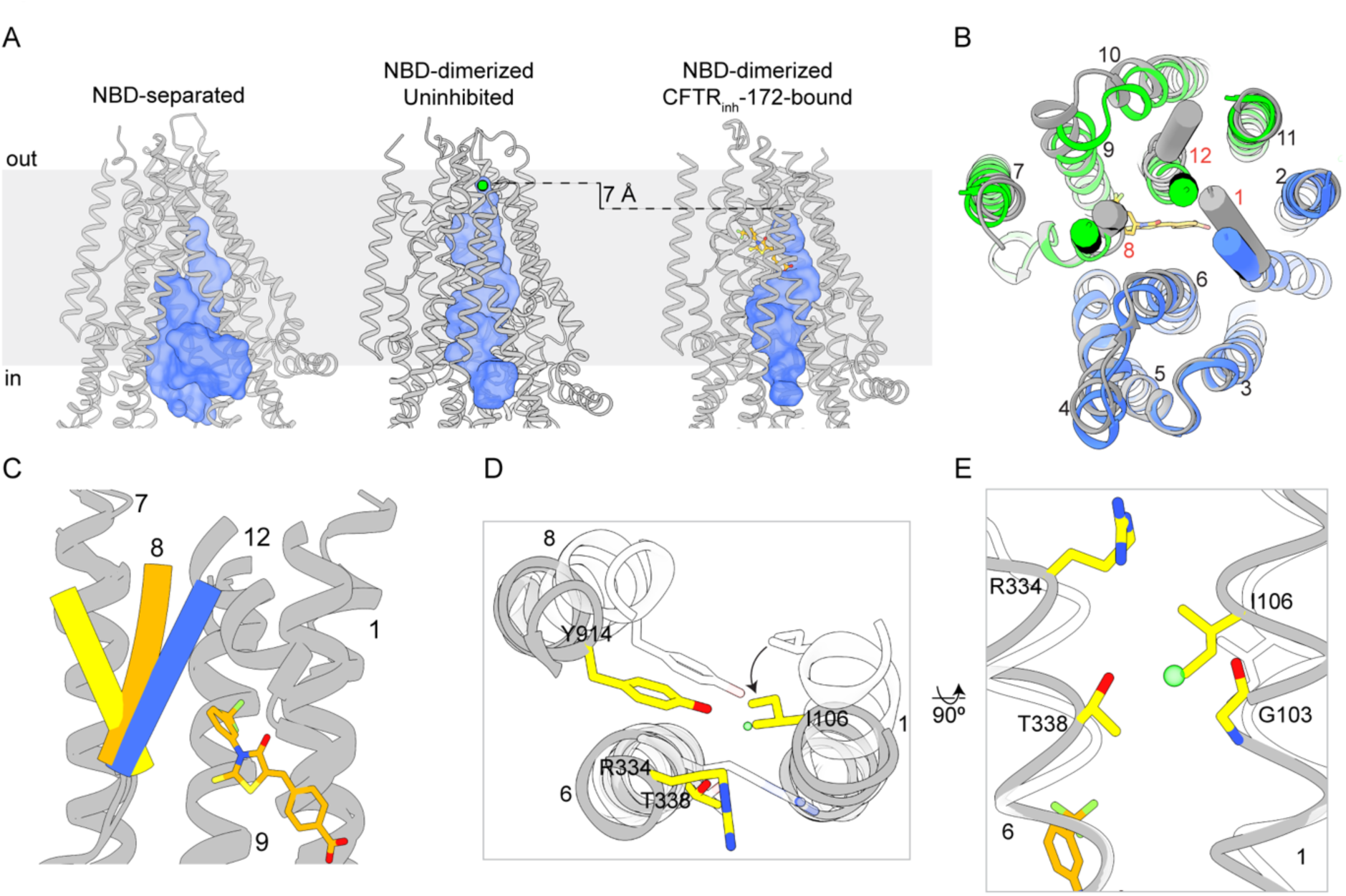
CFTR_inh_-172 stabilizes a closed conformation. (**A**) Comparison of the pore (shown as a blue surface) in NBD-separated (PDB 5UAK), NBD-dimerized, uninhibited (PDB 7SVD) and NBD-dimerized, CFTR_inh_-172-bound CFTR, as defined by a spherical probe with a radius of 1.7 Å. Chloride is modeled as a green sphere in the NBD-dimerized, uninhibited structure. CFTR_inh_-172 is shown as an orange ball-and-stick model. (**B**) Overlay of the CFTR_inh_-172-bound structure (blue and green) with the uninhibited structure (gray). TMs 1, 8, and 12 are shown as cylinders. (**C**) CFTR_inh_-172-stabilized position of TM8 (orange) compared with that observed in the uninhibited structure (blue) and NBD-separated structure (yellow). (**D**) Extracellular view of the local superposition of the helices and residues comprising the selectivity filter in the CFTR_inh_-172-bound (gray/yellow) and uninhibited (transparent white) structures. The chloride ion observed in the uninhibited structure is shown as a green sphere. (**E**) Local superposition of TMs 1 and 6 in the CFTR_inh_-172-bound (gray/yellow) and uninhibited (transparent white) structures. The chloride ion observed in the uninhibited structure is shown as a green sphere.

In the presence of CFTR_inh_-172, CFTR adopts a conformation distinct from either structure (Figure 3A). The NBDs form a dimer similar to that observed in the uninhibited structures of CFTR (E1371Q) (25, 50). The TMDs undergo global rigid-body movements towards each other, but TMs 1, 8 and 12 are positioned differently (Figure 3B). Local structural superposition shows that the extracellular segment of TM8 is stabilized in a conformation intermediate between the NBD-separated and -dimerized conformations (Figure 3C). Furthermore, the anion selectivity filter (see accompanying paper) collapses, as TM1 undergoes a ∼5° rotation that places the side chain of I106 at the position of the chloride ion (Figure 3D-E). This repositioning of the TMs also leads to a complete closure of the lateral exit between TMs 1 and 6 that connects the selectivity filter to the extracellular space (Figure 3E).

### CFTR_inh_-172 allosterically inhibits ATP turnover

The structural analysis clearly shows that CFTR_inh_-172 binds and occludes the pore (Figure 3). However, previous electrophysiological measurements indicate that it does not act as a simple open-channel blocker (45). As CFTR gating is coupled to ATP hydrolysis, we tested if CFTR_inh_-172 changes the ATP turnover rate (Figure 4A). We found that the presence of 10 µM CFTR_inh_-172 decreased saturating ATP turnover (*k_cat_*) by approximately fourfold, from 22.0 ± 2.2 to 5.2 ± 1.0 ATP/protein/min (Figure 4A). The Michaelis–Menten constant (*K_m_*) for ATP was not significantly changed (Figure 4A).

**Figure 4.**
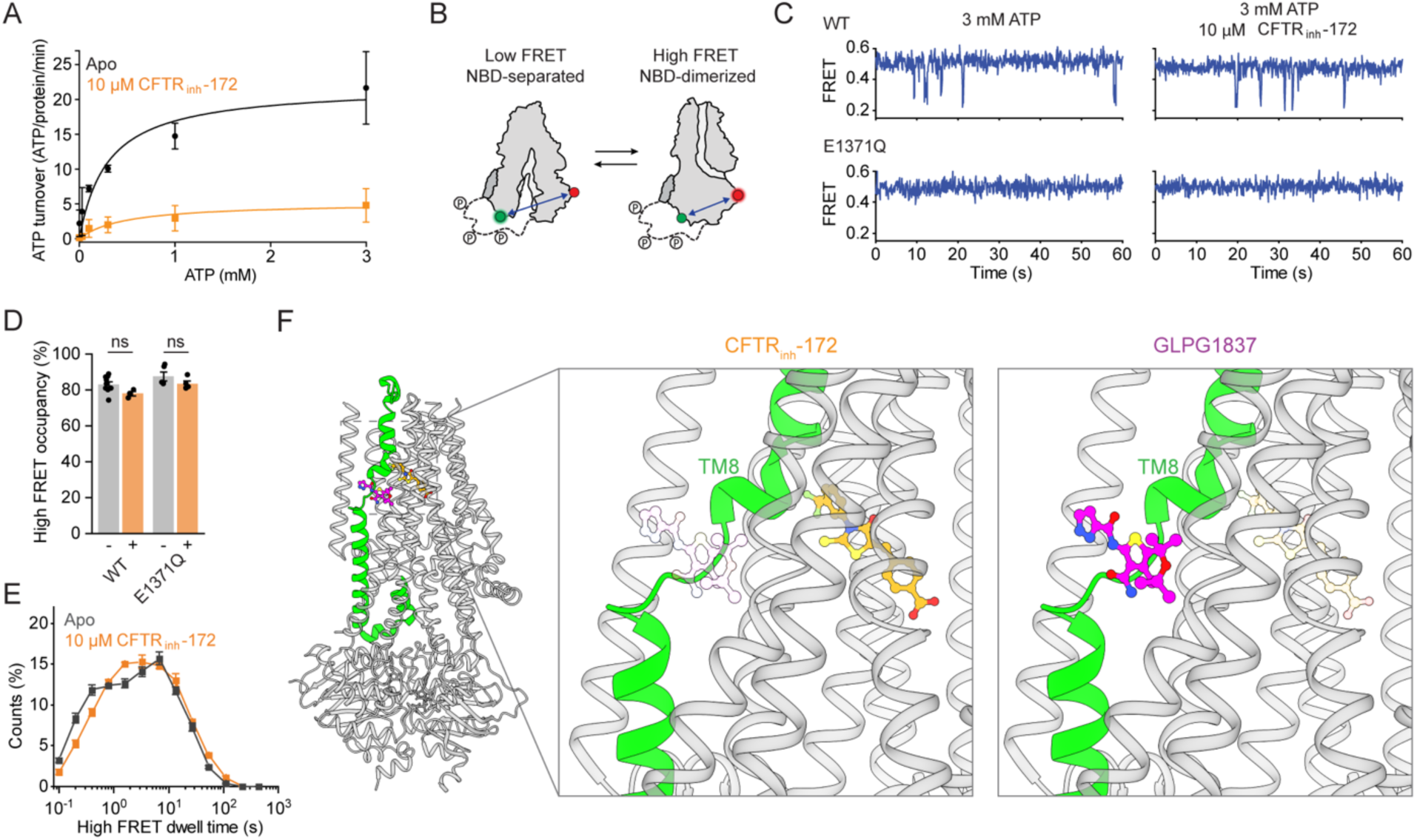
Allosteric inhibition of ATP-dependent gating. (**A**) Effect of CFTR_inh_-172 on steady-state ATP hydrolysis by PKA-phosphorylated WT CFTR. Data represent means and standard errors for 3 (without CFTR_inh_-172) or 4 (with CFTR_inh_-172) measurements and are fitted with the Michaelis-Menten equation. Without CFTR_inh_-172, the *K_m_* for ATP was 0.3 ± 0.1 mM and *k_cat_* was 22 ± 3 ATP/protein/minute. With CFTR_inh_-172, the *K_m_* was 0.47 ± 0.29 mM and *k_cat_* was 5.2 ± 1.0 ATP/protein/minute. (**B**) Schematic of individual CFTR molecules labelled for smFRET imaging. Green and red circles indicate fluorophore positions. (**C**) Example smFRET traces for PKA-phosphorylated WT CFTR and CFTR (E1371Q) with and without CFTR_inh_-172. (**D**) Effects of CFTR_inh_-172 on probabilities of dimerization for WT CFTR and CFTR (E1371Q). Data represent means and standard errors for 9 (WT, without CFTR_inh_-172), 3 (WT, with CFTR_inh_-172), 5 (E1371Q, without CFTR_inh_-172), or 4 (E1371Q, with CFTR_inh_-172) measurements. Statistical significance was tested by one-way analysis of variance (ns, not significant). (**E**) Dwell-time distributions for NBD-dimerization for WT CFTR with and without 10 µM CFTR_inh_-172. Data represent means and standard errors for 8 (without CFTR_inh_-172) or 3 (with CFTR_inh_-172) measurements. (**F**) Cartoon representation of CFTR bound to CFTR_inh_-172 (left). TM 8 is colored green. The potentiator GLPG1837 (colored magenta) was overlayed onto the CFTR_inh_-172-bound structure. Closeup views of the CFTR_inh_-172 (middle) and GLPG1837 (right) binding sites. In each view the other compound is shown as transparent sticks to emphasize their relative positions.

To assess the mechanism by which ATP hydrolysis is inhibited by CFTR_inh_-172, we used a recently established smFRET assay, which reports on the conformational state of CFTR’s NBDs (38). In this assay, position 388 in NBD1 and position 1435 in NBD2 were labeled with donor and acceptor fluorophores (Figure 4B). Conformational isomerizations of individual CFTR molecules were monitored as transitions between a low FRET efficiency (0.25 ± 0.01) NBD-separated state and a high FRET efficiency (0.49 ± 0.02) NBD-dimerized state. As we have previously reported (38), at a saturating (3 mM) ATP concentration, WT CFTR predominantly occupies NBD-dimerized conformations with brief excursions to the NBD-separated state (Figure 4C). The presence of 10 µM CFTR_inh_-172 did not significantly affect the probability of NBD dimerization or the dwell time of the NBD-dimerized state for WT CFTR (Figure 4D-E). CFTR_inh_-172 also did not affect the conformational dynamics of CFTR (E1371Q), which remained constitutively NBD-dimerized (Figure 4C-E). These data indicate that CFTR_inh_-172 does not prevent NBD dimerization, but rather slows progression through the gating cycle while the NBDs are dimerized.

These observations lead us to consider a possible mechanism for inhibition of ATP hydrolysis. Our recent study showed that conformational changes within NBD-dimerized CFTR governed by ATP turnover are required for chloride conductance (38). Potentiators Ivacaftor and GLPG1837 enhance channel activity by increasing pore opening while the NBDs are dimerized. Additionally, the potentiators increase ATP turnover (38). In comparing the structure of CFTR (E1371Q) bound to GLPG1837 with that bound to CFTR_inh_-172, we observe that the CFTR_inh_-172 site is located along the pore-lining side of the TM8 hinge region, in direct juxtaposition to the potentiator binding site (Figure 4F). As TM8 links ATP hydrolysis and pore opening (36, 51), we propose that CFTR_inh_-172 inhibits ATP turnover via an allosteric mechanism involving TM8, similar in nature but opposite in effect to that of the potentiators.

## DISCUSSION

Typically, ion channel inhibitors are classified into two categories: pore blockers that bind within the ion conduction pathway to occlude the pore and gating inhibitors that impair channel opening by stabilizing the closed state (52). However, CFTR_inh_-172 presents a perplexing case, as it has been shown to interact with residues within the pore and also impair gating (44, 45). Through cryo-EM and smFRET analyses, we now have a structural understanding of CFTR_inh_-172 inhibition that reconciles earlier findings. The binding site of CFTR_inh_-172 is located within the pore, nestled in a cavity lined by R347, a residue whose substitution significantly reduces the affinity for CFTR_inh_-172 (44). Whereas the principal mechanism of CFTR_inh_-172 involves steric occlusion of the pore, we also observe local conformational changes of TMs 1, 8, and 12. These changes cause a collapse of the chloride selectivity filter and the extracellular exit. Based on kinetic analysis Hwang and colleagues had predicted that CFTR_inh_-172 induces conformational change of CFTR (46). The nature of this change is now revealed at a molecular level. Additionally, we discovered that CFTR_inh_-172 inhibits ATP hydrolysis through an allosteric mechanism similar to that of potentiators, but with an opposite functional effect. These observations corroborate the hypothesis that conformational shifts in TM8 link ATP hydrolysis at the NBDs with the state of the pore.

Electrophysiological measurements in our laboratory (Figure 1) and in other studies (46) demonstrate that CFTR_inh_-172 inhibits WT CFTR and several hydrolysis-deficient variants to similar extents. Although it is theoretically possible that this molecule inhibits WT CFTR and each variant through different mechanisms, it is more likely that the mechanism of action is similar. Indeed, the structure determined with CFTR (E1371Q) is entirely consistent with earlier SAR studies (5, 48). Substitutions at the structurally identified binding site made in the WT CFTR background reduced the potency of CFTR_inh_-172 (44) (Figure 2E). Additionally, smFRET studies reveal that CFTR_inh_-172 does not affect the NBD isomerization in WT CFTR or CFTR (E1371Q) (Figure 4C-D). These data strongly suggest that the mode of action revealed in this study represents a general mechanism for CFTR_inh_-172. However, E1371Q substitution stabilizes the NBDs in a canonical dimerized conformation, and it is possible that CFTR_inh_-172 induces local changes at the ATPase site that are obscured in our structure. Further studies will be pursued to identify structural re-arrangements of the WT channel within the NBD-dimerized state, as these play a key role in coupling ATP hydrolysis to channel gating.

Finally, the congruence between structural and functional data not only offers intellectual satisfaction but also opens up new avenues for enhancing the potency and specificity of CFTR_inh_-172. Specifically, CFTR interacts with Ring C of CFTR_inh_-172 primarily through the K95 salt-bridge and an edge-to-face π-stacking interaction with W1145. It is possible that analogs of CFTR_inh_-172, with modifications on Ring C that establish additional interactions with nearby polar residues on CFTR will have enhanced potency and specificity.

## METHODS

### Cell culture

Sf9 cells (Gibco, catalogue number 11496015, lot number 1670337) were grown at 27 °C in Sf-900 II SFM medium (Gibco) supplemented with 5% (v/v) fetal bovine serum (FBS) and 1% (v/v) antibiotic-antimycotic (Gibco). HEK293S GnTI^-^ cells (ATCC CRL-3022, lot number 62430067) were cultured at 37 °C in Freestyle 293 medium (Gibco) supplemented with 2% (v/v) FBS and 1% (v/v) antibiotic-antimycotic. CHO-K1 cells (ATCC CCL-61, lot number 70014310) were cultured at 37 °C in DMEM F-12 medium (ATCC) supplemented with 10% (v/v) FBS and 1% (v/v) GlutaMAX (Gibco).

### Mutagenesis

CFTR mutants were generated using the SPRINP mutagenesis method (**Table S2**) (53). Briefly, mutagenic primers were designed to be complementary to the template plasmid except for the mutated bases and to be 15-45 nucleotides in length. Plasmid containing CFTR cDNA was amplified in separate reactions containing forward or reverse primer. The single-primer products of these reactions were combined and denatured at 95 °C for 5 min. and gradually cooled to 37 °C over the next 5 min. The sample was then digested by DpnI for 4 hours. 5 µL of sample were added to 50 µL of competent XL2Blue cells for transformation and incubated on ice for 30 min. The bacteria were then heat-shocked at 42 °C for 45 seconds and allowed to recover on ice for 2 min. 200 µL of warmed SOC media (Invitrogen) were then added directly to the cells, and the mixture was allowed to shake at 225 RPM in a 37 °C incubator. 200 µL of this mixture was then spread on LB/ampicillin plates and left to incubate at 37 °C overnight. Random colonies were then picked and expanded in LB/ampicillin. Plasmid DNA was then purified (QIAGEN Plasmid Kit) and sequenced (Genewiz).

### Patch-clamp electrophysiology

Chinese hamster ovary (CHO; ATCC CCL-61, lot number 70014310) cells were maintained in DMEM-F12 (ATCC) supplemented with 10% (v/v) heat-inactivated fetal bovine serum (FBS) and 1% GlutaMAX (Gibco) at 37°C. The cells were seeded in 35-mm cell culture dishes (Falcon) 24 hours before transfection. Cells were transiently transfected with BacMam vector encoding C-terminally GFP-fused CFTR, using Lipofectamine 3000 (Invitrogen). 12 hours after transfection, medium was exchanged for DMEM-F12 supplemented with 2% (v/v) FBS and 1% (v/v) GlutaMAX and incubation temperature was reduced to 30 °C. Patch-clamp recording was carried out after an additional 24 hours.

The bath solution was 145 mM NaCl, 2 mM MgCl_2_, 5 mM KCl, 1 mM CaCl_2_, 5 mM glucose, 5 mM HEPES and 20 mM sucrose (pH 7.4 with NaOH). Pipette solution was 140 mM NMDG, 5 mM CaCl_2_, 2 mM MgCl_2_ and 10 mM HEPES (pH 7.4 with HCl). Perfusion solution was 150 mM NMDG, 2 mM MgCl_2_, 1 mM CaCl_2_, 10 mM EGTA and 8 mM Tris (pH 7.4 with HCl).

Recordings were carried out using the inside-out patch configuration with local perfusion at the patch. Recording pipettes were pulled from borosilicate glass (outer diameter 1.5 mm, inner diameter 0.86 mm, Sutter) to 1.5–3.0 MΩ resistance. Currents were recorded at 25°C using an Axopatch 200B amplifier, a Digidata 1550 digitizer and the pClamp software suite (Molecular Devices). Membrane potential was clamped at -30 mV. Current traces reflect inward currents with inverted signatures. Recordings were low-pass-filtered at 1 kHz and digitized at 20 kHz.

For all measurements, CFTR was activated by exposure to PKA (Sigma-Aldrich) and 3 mM ATP. Displayed recordings were low-pass filtered at 100 Hz. Data were analyzed using Clampfit, GraphPad Prism, and OriginPro.

### Protein expression and purification

CFTR constructs were expressed and purified as previously described (54, 55). Bacmids encoding human CFTR fused to a C-terminal PreScission Protease-cleavable green fluorescent protein (GFP) tag were generated in *E. coli* DH10Bac cells (Invitrogen). Recombinant baculovirus was produced and amplified in Sf9 cells. HEK293S GnTl^-^ suspension cells, at a density of 2.0-3.0 × 10^6^ cells/ml, were infected with 10% (v/v) P3 or P4 baculovirus. Protein expression was induced by addition of 10 mM (final concentration) sodium butyrate 12 hours after infection. The cells were cultured at 30°C for an additional 48 hours and then harvested by centrifugation.

Protein samples for cryo-EM were purified as follows. Cells were solubilized for 75 min at 4°C in extraction buffer containing 1-1.25% (w/v) 2,2-didecylpropane-1,3-bis-b-D-maltopyranoside (LMNG), 0.25% (w/v) cholesteryl hemisuccinate (CHS), 200 mM NaCl, 20 mM HEPES (pH 7.5 with NaOH), 2mM MgCl_2_, 10 μM dithiothreitol (DTT), 20% (v/v) glycerol, 2 mM ATP, 1 μg mL^-1^ pepstatin A, 1μg mL^-1^aprotinin, 100μg mL^-1^ soy trypsin inhibitor, 1mM benzamidine, 1mM phenylmethylsulfonyl fluoride (PMSF) and 3 μg mL^-1^ DNase I. Lysate was clarified by centrifugation at 75,000*g* for 45 min at 4°C, and mixed with NHS-activated Sepharose 4 Fast Flow resin (GE Healthcare) conjugated with GFP nanobody, which had been pre-equilibrated in extraction buffer. After 2 hours, the resin was packed into a chromatography column and was washed with buffer containing 0.06% (w/v) digitonin, 200 mM NaCl, 20 mM HEPES (pH 7.5 with NaOH), 2 mM ATP, and 2 mM MgCl_2_. The resin was then incubated for 2 hours at 4°C with 0.35 mg mL^-1^ PreScission Protease to cleave off the GFP tag. The PreScission Protease was removed by dripping eluate through Glutathione Sepharose 4B resin (Cytiva). The protein was then concentrated to yield ∼1mL of sample, which was then phosphorylated by protein kinase A (PKA). Finally, protein samples were purified by size-exclusion chromatography at 4°C using a Superose 6 10/300 GL column (GE Healthcare), equilibrated with a buffer containing 0.03% (w/v) digitonin, 200 mM NaCl, 20 mM HEPES (pH 7.5 with NaOH), 2 mM ATP, and 2 mM MgCl_2_. Peak fractions were pooled and concentrated.

Samples for ATP hydrolysis assays were purified using the same protocol but in a buffer containing KCl, rather than NaCl. The sample used for single-molecule FRET imaging had the following substitutions: C76L, C128S, C225S, C276S, C343S, T388C, C491S, C592M, C647S, C832S, C866S, C1344S, C1355S, C1395S, C1400S, C1410S, S1435C and C1458S. The purification protocol was adjusted as described in (38). For proteoliposome reconstitution, the size-exculsion, chromatography buffer contained glyco-diosgenin (GDN) instead of digitonin.

### EM data acquisition and processing

Immediately following size-exclusion chromatography, the CFTR (E1371Q) sample was concentrated to 5 mg/mL (32 µM) and incubated with 8 mM ATP, 10 mM MgCl_2_, and 100 µM CFTR_inh_-172 on ice for 30 min. 3 mM fluorinated Fos-choline-8 was added to the samples directly before application onto glow-discharged Quantifoil R0.6/1 300 mesh Cu grids. Samples were then vitrified using a Vitrobot Mark IV (FEI).

Cryo-EM images were collected in a super-resolution mode on a 300 kV Titan Krios (FEI) equipped with a K3 Summit detector (Gatan) using SerialEM (**Table S1**). Images were corrected for gain reference and binned by 2. Drift correction was performed using MotionCor2 (56). Contrast transfer function (CTF) estimation was performed using GCTF (57). GCTF values were used for further processing steps. Particles were picked using Gautomatch (https://www.mrc-lmb.cam.ac.uk/kzhang/). All subsequent steps of map reconstruction and resolution estimation were carried out using RELION 3.1 (58) (**Figure S1**).

After the first round of 3D classification, 3D refinement was performed, followed by an additional round of 3D classification without alignment and 3D refinement. The resulting dataset was then put through three rounds of CTF refinement, the first and third of which estimated anisotropic magnification. The dataset was then put through another round of 3D refinement, after which it was polished and refined again.

### Model building and refinement

Initial protein models were built by fitting the published structure of the NBD-dimerized CFTR (E1371Q) (PDB: 6MSM) into the cryo-EM map using Coot (59). The model was then adjusted based on the cryo-EM density. CFTR_inh_-172 was built into the density and refined in PHENIX (60) using restraints generated by the Global Phasing web server (grade.globalphasing.org). MolProbity (61) was used for geometry validation.

### ATP hydrolysis measurements

Steady-state ATP hydrolysis was measured using an NADH-coupled assay (62). The assay buffer contained 50 mM HEPES (pH 8.0 with KOH), 150 mM KCl, 2 mM MgCl_2_, 2 mM DTT, 0.06% (w/v) digitonin, 60 µg/ml pyruvate kinase (Roche), 32 µg/ml lactate dehydrogenase (Roche), 9 mM phosphoenolpyruvate, 150 µM NADH, and 200 nM CFTR. Aliquots of 27 µL were distributed into a Corning 384-well Black/Clear Flat Bottom Polystyrene NBS Microplate. The reactions were initiated by the addition of ATP. The rate of fluorescence depletion was monitored at λ_ex_ = 340 nm and λ_em_ = 445 nm at 28 °C with an Infinite M1000 microplate reader (Tecan). ATP turnover was then determined with an NADH standard curve.

### Proteoliposome reconstitution and planar bilayer recording

The lipids 1,2-dioleoyl-*sn*-glycero-3-phosphoethanolamine (DOPE), 1-palmitoyl-2-oleyl-*sn*-glycero-3-phosphocholine (POPC), and 1-palmitoyl-2-oleoyl-*sn*-glycero-3-phospho-L-serine (POPS) were mixed at a 2:1:1 (w/w/w) ratio and resuspended by sonication in buffer containing 200 mM NaCl, 20 mM HEPES (pH 7.2 with NaOH) and 2 mM MgCl_2_ to a final lipid concentration of 20 mg/ml. 2% (w/v) GDN was added and the mixture was incubated for 1 hour at 25 °C. CFTR was mixed with the lipids at a protein-to-lipid ratio of 1:250 (w/w) and incubated at 4 °C for 2 hours. 14 mg/ml methylated beta-cyclodextrin was added to the mixture. After 4 hours an equivalent amount of methylated beta-cyclodextrin was added to the mixture. This was performed for a total of four additions. Proteoliposomes were pelleted by centrifugation at 150,000*g* for 45 min at 4 °C and resuspended in buffer containing 200 mM NaCl, 20 mM HEPES (pH 7.2 with NaOH) and 2 mM MgCl_2_.

Synthetic planar lipid bilayers were made from a lipid mixture containing DOPE, POPC, and POPS at a 2:1:1 (w/w/w) ratio. Proteoliposomes containing PKA-phosphorylated CFTR were fused with the bilayers. Currents were recorded at 25 °C in a symmetric buffer containing 150 mM NaCl, 2 mM MgCl_2_, 20 mM HEPES (pH 7.2 with NaOH), and 3 mM ATP. Voltage was clamped at -150 mV with an Axopatch 200B amplifier (Molecular Devices). Currents were low-pass filtered at 1 kHz, digitized at 20 kHz with a Digidata 1440A digitizer and recorded using the pCLAMP software suite (Molecular devices). Recordings were further low-pass filtered at 100 Hz. Data were analyzed with Clampfit, GraphPad Prism and OriginPro.

### Single-molecule fluorescence imaging and FRET data analysis

Imaging and analysis were carried out as outlined in (38, 63). In brief, PKA-phosphorylated CFTR was immobilized within PEG-passivated microfluidic chambers via a streptavidin–biotin–tris-(NTA-Ni^2+^) bridge. Experiments were performed at 25 °C in buffer containing 0.06% (w/v) digitonin, 150 mM NaCl, 2 mM MgCl_2_, 20 mM HEPES (pH 7.2 with NaOH), 2 mM protocatechuic acid and 50 nM protocatechuate-3,4-dioxygenase.

Imaging was carried out with a custom-built wide-field, prism-based total internal reflection fluorescence microscope. Donor (LD555) fluorophores were excited with an evanescent wave generated using a 532-nm laser (Opus, Laser Quantum). Fluorescence emitted from donor (LD555) and acceptor (LD655) fluorophores was collected with a 1.27 NA 60× water-immersion objective (Nikon), spectrally resolved using a T635lpxr dichroic (Chroma), and imaged onto two Fusion sCMOS cameras (Hamamatsu). The integration period of imaging was 100 ms.

Analysis of fluorescence data was performed using the SPARTAN analysis software in MATLAB (64). Single-molecule FRET trajectories were calculated as *E_FRET_* = *I_A_*/(*I_A_* + *I_D_*), where *I_A_* and *I_D_* are the emitted acceptor and donor fluorescence intensities, respectively. The following pre-established criteria were applied to select FRET trajectories for analysis: single-step donor photobleaching; a signal-to-noise ratio above 8; fewer than 4 donor-blinking events; FRET efficiency below 0.8; and FRET efficiency above baseline for at least 50 frames. The segmental k-means algorithm (65) was used to idealize trajectories to a model containing two non-zero-FRET states with FRET efficiencies of 0.25 and 0.48. Data were further analyzed with OriginPro.

---

**Figure S1.**
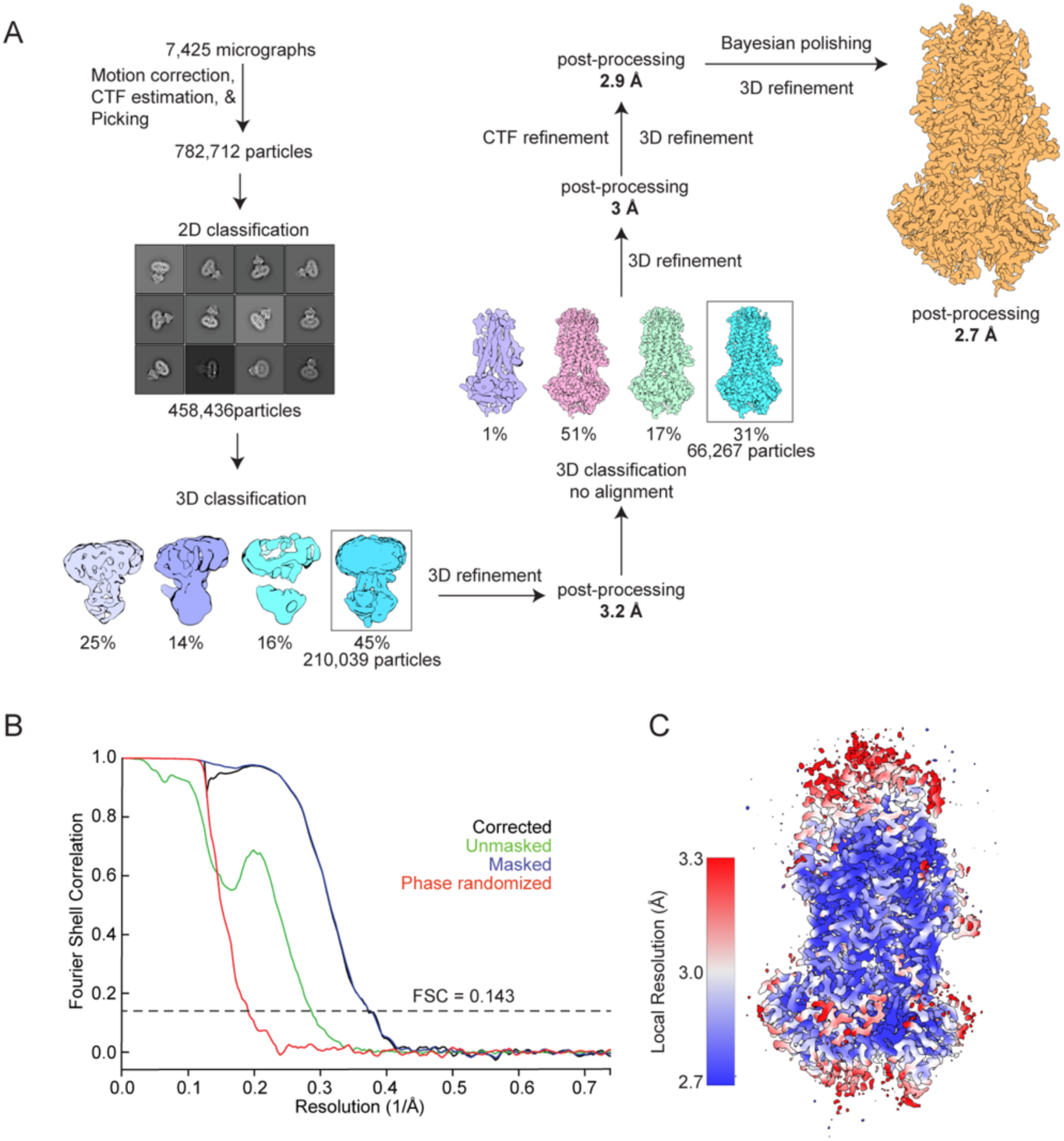
Cryo-EM analysis of the CFTR/CFTR_inh_-172 complex and quality of the reconstruction. (**A**) Image processing procedure. (**B**) Fourier shell correlation curves of the final map. (**C**) Local resolution estimation of the final map.

**Figure S2.**
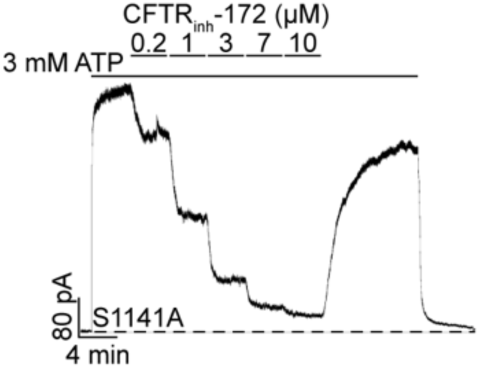
CFTR_inh_-172 titration onto S1141A CFTR. Example macroscopic current traces showing titration of CFTR_inh_-172 onto S1141A CFTR in inside-out excised patches. CFTR was fully phosphorylated by PKA in the presence of 3 mM ATP before CFTR_inh_-172 titration.

**Table S1.** Cryo-EM data collection, refinement, and validation statistics.

|  |  |
| --- | --- |
|  | CFTR (E1371Q)/CFTR<sub>inh</sub>-172 +ATP/Mg<sup>2+</sup><br>(EMDB-42101)<br>(PDB 8UBR) |
| **Data collection and processing** |  |
| Magnification | 105,000 |
| Voltage (kV) | 300 |
| Electron exposure (e<sup>-</sup>/Å<sup>2</sup>) | 65.6 |
| Defocus range (μm) | 0.5-2.5 |
| Pixel size (Å) | 0.676 |
| Symmetry imposed | C1 |
| Initial particle images (no.) | 782,712 |
| Final particle images (no.) | 66,267 |
| Map resolution (Å) | 2.70 |
| FSC threshold | 0.143 |
| Map resolution range (Å) | 2.7-3.8 |
| **Refinement** |  |
| Initial model used (PDB code) | 6MSM |
| Model resolution (Å) | 2.6 |
| FSC threshold | 0.143 |
| Map sharpening *B* factor (Å<sup>2</sup>) | -43.286 |
| Model composition |  |
| Non-hydrogen atoms | 9440 |
| Protein residues | 1156 |
| Ligands | 9 |
| *B* factors (Å<sup>2</sup>) |  |
| Protein | 98 |
| Ligand (inhibitor) | 67 |
| R.m.s. deviations |  |
| Bond lengths (Å) | 0.070 |
| Bond angles (°) | 0.870 |
| Validation |  |
| MolProbity score | 1.17 |
| Clashscore | 3.811 |
| Poor rotamers (%) | 0.1 |
| Ramachandran plot |  |
| Favored (%) | 98.25 |
| Allowed (%) | 1.75 |
| Disallowed (%) | 0 |

**Table S2.**
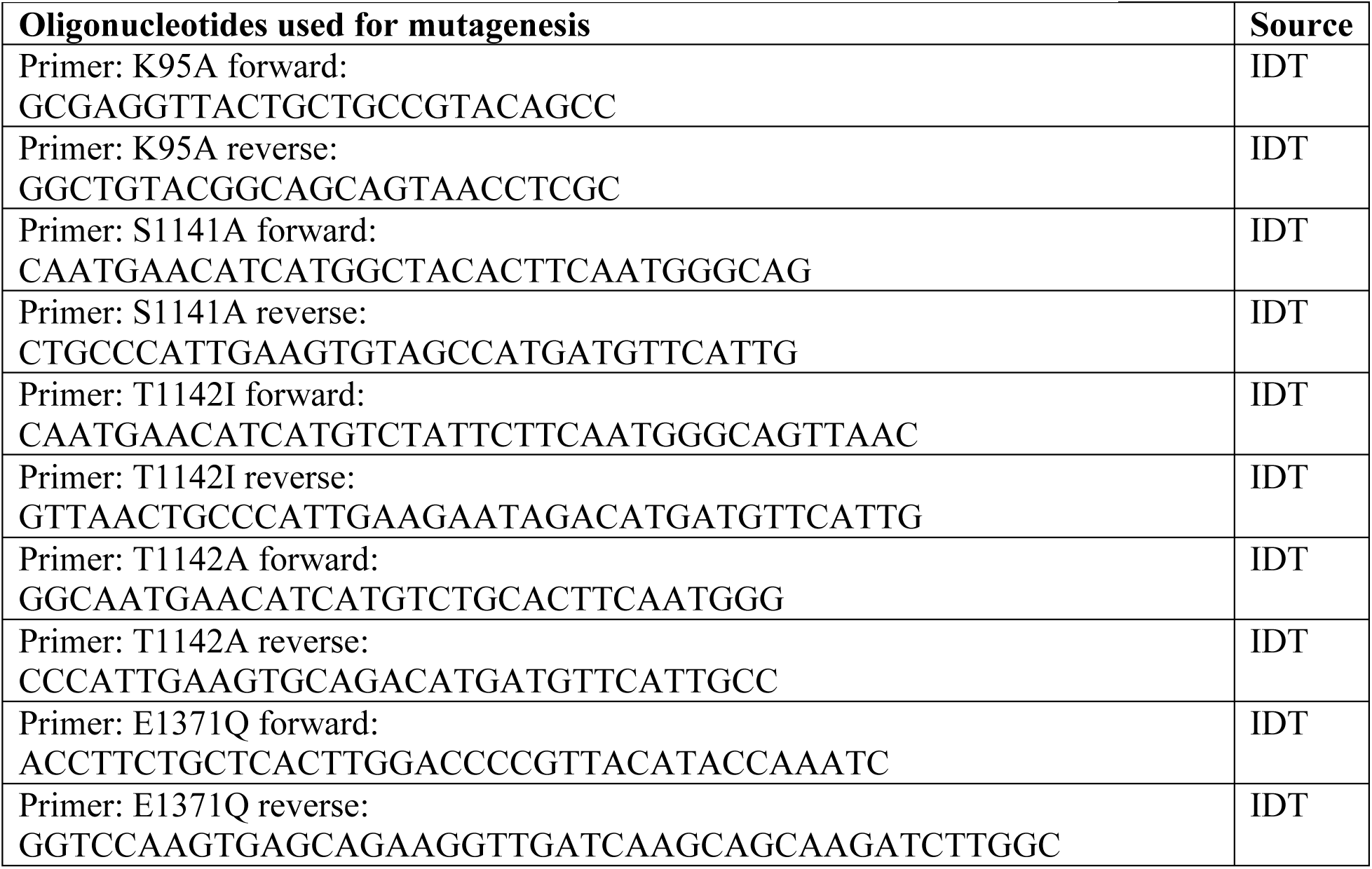
List of oligonucleotides used in this study.

## Acknowledgements

We acknowledge the support from Mark Ebrahim, Johanna Sotiris, and Honkit Ng at Rockefeller’s Evelyn Gruss Lipper Cryo-Electron Microscopy Resource Center in collecting electron microscopy data, and the assistance from the Single-Molecule Imaging Center at St. Jude Children’s Research Hospital in single-molecule fluorescence imaging. We thank members of the Chen and Mackinnon laboratories for helpful discussions and Dr. L. Csanády for comments on the manuscript This work was supported by HHMI to J.C., The Cystic Fibrosis Foundation Therapeutics to K.F., the National Institutes of Health to S.C.B.(GM079238) and P.Y (Training grant, T32GM007739).

## Author contributions

P.Y. and K.F determined the cryo-EM structure; P.Y. performed ATP turnover assays and patch-clamp experiments presented in Figure 2E; J.L. performed electrophysiology experiments presented in Figure 1 and the single-molecule FRET experiments. All authors contributed to writing the manuscript. S.C.B. and J.C. oversaw the project.

## Data and materials availability

The cryo-EM map of CFTR_inh_-172-bound CFTR has been deposited in the Electron Microscopy Data Bank under the accession code EMD-42101. The corresponding atomic model has been deposited in the Protein Data Bank under accession code 8UBR. All other data and information are available in the main text or the supplementary materials.

## References

1. D. C. Gadsby, P. Vergani, L. Csanády, The ABC protein turned chloride channel whose failure causes cystic fibrosis. Nature 440, 477–483 (2006).

2. M. P. Anderson, et al., Demonstration that CFTR is a chloride channel by alteration of its anion selectivity. Science 253, 202–205 (1991).

3. B. Yang, N. D. Sonawane, D. Zhao, S. Somlo, A. S. Verkman, Small-molecule CFTR inhibitors slow cyst growth in polycystic kidney disease. J. Am. Soc. Nephrol. JASN 19, 1300–1310 (2008).

4. J. R. Thiagarajah, T. Broadbent, E. Hsieh, A. S. Verkman, Prevention of toxin-induced intestinal ion and fluid secretion by a small-molecule CFTR inhibitor. Gastroenterology 126, 511–519 (2004).

5. T. Ma, et al., Thiazolidinone CFTR inhibitor identified by high-throughput screening blocks cholera toxin–induced intestinal fluid secretion. J. Clin. Invest. 110, 1651–1658 (2002).

6. D. S. Snyder, L. Tradtrantip, C. Yao, M. J. Kurth, A. S. Verkman, Potent, Metabolically Stable Benzopyrimido-pyrrolo-oxazine-dione (BPO) CFTR Inhibitors for Polycystic Kidney Disease. J. Med. Chem. 54, 5468–5477 (2011).

7. F. A. Belibi, et al., Cyclic AMP promotes growth and secretion in human polycystic kidney epithelial cells1 1See Editorial by Torres, p. 1283. Kidney Int. 66, 964–973 (2004).

8. C. J. Davidow, R. L. Maser, L. A. Rome, J. P. Calvet, J. J. Grantham, The cystic fibrosis transmembrane conductance regulator mediates transepithelial fluid secretion by human autosomal dominant polycystic kidney disease epithelium in vitro. Kidney Int. 50, 208–218 (1996).

9. J. R. Thiagarajah, M. Donowitz, A. S. Verkman, Secretory diarrhoea: mechanisms and emerging therapies. Nat. Rev. Gastroenterol. Hepatol. 12, 446–457 (2015).

10. C. Moon, et al., Drug-induced secretory diarrhea: A role for CFTR. Pharmacol. Res. 102, 107–112 (2015).

11. A. C. Chao, et al., Activation of intestinal CFTR Cl-channel by heat-stable enterotoxin and guanylin via cAMP-dependent protein kinase. EMBO J. 13, 1065–1072 (1994).

12. V. E. Torres, P. C. Harris, Strategies Targeting cAMP Signaling in the Treatment of Polycystic Kidney Disease. J. Am. Soc. Nephrol. 25, 18 (2014).

13. K. Hanaoka, W. B. Guggino, cAMP regulates cell proliferation and cyst formation in autosomal polycystic kidney disease cells. J. Am. Soc. Nephrol. JASN 11, 1179–1187 (2000).

14. C. S. Hyun, G. A. Kimmich, Effect of cholera toxin on cAMP levels and Na+ influx in isolated intestinal epithelial cells. Am. J. Physiol. 243, C107–115 (1982).

15. J. K. Scholz, et al., Loss of Polycystin-1 causes cAMP-dependent switch from tubule to cyst formation. iScience 25, 104359 (2022).

16. D. M. Gill, Involvement of nicotinamide adenine dinucleotide in the action of cholera toxin in vitro. Proc. Natl. Acad. Sci. 72, 2064–2068 (1975).

17. D. M. Gill, R. Meren, ADP-ribosylation of membrane proteins catalyzed by cholera toxin: basis of the activation of adenylate cyclase. Proc. Natl. Acad. Sci. 75, 3050–3054 (1978).

18. D. Cassel, T. Pfeuffer, Mechanism of cholera toxin action: Covalent modification of the guanyl nucleotide-binding protein of the adenylate cyclase system. Proc. Natl. Acad. Sci. 75, 2669–2673 (1978).

19. V. Patel, R. Chowdhury, P. Igarashi, Advances in the Pathogenesis and Treatment of Polycystic Kidney Disease. Curr. Opin. Nephrol. Hypertens. 18, 99 (2009).

20. J. B. Harris, R. C. LaRocque, F. Qadri, E. T. Ryan, S. B. Calderwood, Cholera. The Lancet 379, 2466–2476 (2012).

21. S. Rajak, et al., Pharmacological inhibition of CFTR attenuates nonalcoholic steatohepatitis (NASH) progression in mice. Biochim. Biophys. Acta Mol. Basis Dis. 1869, 166662 (2023).

22. L. Csanády, P. Vergani, D. C. Gadsby, STRUCTURE, GATING, AND REGULATION OF THE CFTR ANION CHANNEL. Physiol. Rev. 99 (2019).

23. T.-C. Hwang, et al., Structural mechanisms of CFTR function and dysfunction. J. Gen. Physiol. 150, 539–570 (2018).

24. S. H. Cheng, et al., Phosphorylation of the R domain by cAMP-dependent protein kinase regulates the CFTR chloride channel. Cell 66, 1027–1036 (1991).

25. Z. Zhang, F. Liu, J. Chen, Conformational Changes of CFTR upon Phosphorylation and ATP Binding. Cell 170, 483–491.e8 (2017).

26. L. Csanády, P. Vergani, D. C. Gadsby, Strict coupling between CFTR’s catalytic cycle and gating of its Cl-ion pore revealed by distributions of open channel burst durations. Proc. Natl. Acad. Sci. 107, 1241–1246 (2010).

27. S. Hadida, et al., Discovery of N-(2,4-Di-tert-butyl-5-hydroxyphenyl)-4-oxo-1,4-dihydroquinoline-3-carboxamide (VX-770, Ivacaftor), a Potent and Orally Bioavailable CFTR Potentiator. J. Med. Chem. 57, 9776–9795 (2014).

28. S. E. Van der Plas, et al., Discovery of N-(3-Carbamoyl-5,5,7,7-tetramethyl-5,7-dihydro-4H-thieno[2,3-c]pyran-2-yl)-lH-pyrazole-5-carboxamide (GLPG1837), a Novel Potentiator Which Can Open Class III Mutant Cystic Fibrosis Transmembrane Conductance Regulator (CFTR) Channels to a High Extent. J. Med. Chem. 61, 1425–1435 (2018).

29. J. L. Taylor-Cousar, et al., Tezacaftor–Ivacaftor in Patients with Cystic Fibrosis Homozygous for Phe508del. N. Engl. J. Med. 377, 2013–2023 (2017).

30. C. E. Wainwright, et al., Lumacaftor–Ivacaftor in Patients with Cystic Fibrosis Homozygous for Phe508del CFTR. N. Engl. J. Med. 373, 220–231 (2015).

31. B. W. Ramsey, et al., A CFTR Potentiator in Patients with Cystic Fibrosis and the G551D Mutation. N. Engl. J. Med. 365, 1663–1672 (2011).

32. D. Keating, et al., VX-445–Tezacaftor–Ivacaftor in Patients with Cystic Fibrosis and One or Two Phe508del Alleles. N. Engl. J. Med. 379, 1612–1620 (2018).

33. J. C. Davies, et al., VX-659–Tezacaftor–Ivacaftor in Patients with Cystic Fibrosis and One or Two Phe508del Alleles. N. Engl. J. Med. 379, 1599–1611 (2018).

34. K.-Y. Jih, T.-C. Hwang, Vx-770 potentiates CFTR function by promoting decoupling between the gating cycle and ATP hydrolysis cycle. Proc. Natl. Acad. Sci. 110, 4404–4409 (2013).

35. F. Van Goor, et al., Rescue of CF airway epithelial cell function in vitro by a CFTR potentiator, VX-770. *Proc. Natl. Acad. Sci.* **106**, 18825–18830 (2009).

36. F. Liu, et al., Structural identification of a hotspot on CFTR for potentiation. Science 364, 1184–1188 (2019).

37. K. Fiedorczuk, J. Chen, Molecular structures reveal synergistic rescue of Δ508 CFTR by Trikafta modulators. Science 378, 284–290 (2022).

38. J. Levring, et al., CFTR function, pathology and pharmacology at single-molecule resolution. Nature, 1–9 (2023).

39. J. Bryce, C. Boschi-Pinto, K. Shibuya, R. E. Black, WHO estimates of the causes of death in children. The Lancet 365, 1147–1152 (2005).

40. F. Rivera-Chávez, B. T. Meader, S. Akosman, V. Koprivica, J. J. Mekalanos, A Potent Inhibitor of the Cystic Fibrosis Transmembrane Conductance Regulator Blocks Disease and Morbidity Due to Toxigenic Vibrio cholerae. Toxins 14, 225 (2022).

41. G. de Wilde, et al., Identification of GLPG/ABBV-2737, a Novel Class of Corrector, Which Exerts Functional Synergy With Other CFTR Modulators. Front. Pharmacol. 10 (2019).

42. C. Muanprasat, et al., Discovery of glycine hydrazide pore-occluding CFTR inhibitors: mechanism, structure-activity analysis, and in vivo efficacy. J. Gen. Physiol. 124, 125–137 (2004).

43. D. S. Snyder, L. Tradtrantip, C. Yao, M. J. Kurth, A. S. Verkman, Potent, Metabolically Stable Benzopyrimido-pyrrolo-oxazine-dione (BPO) CFTR Inhibitors for Polycystic Kidney Disease. J. Med. Chem. 54, 5468–5477 (2011).

44. E. Caci, et al., Evidence for direct CFTR inhibition by CFTR(inh)-172 based on Arg347 mutagenesis. Biochem. J. 413, 135–142 (2008).

45. A. Taddei, et al., Altered channel gating mechanism for CFTR inhibition by a high-affinity thiazolidinone blocker. FEBS Lett. 558, 52–56 (2004).

46. Z. Kopeikin, Y. Sohma, M. Li, T.-C. Hwang, On the mechanism of CFTR inhibition by a thiazolidinone derivative. J. Gen. Physiol. 136, 659–671 (2010).

47. P. Vergani, S. W. Lockless, A. C. Nairn, D. C. Gadsby, CFTR channel opening by ATP-driven tight dimerization of its nucleotide-binding domains. Nature 433, 876–880 (2005).

48. N. D. Sonawane, A. S. Verkman, Thiazolidinone CFTR inhibitors with improved water solubility identified by structure-activity analysis. Bioorg. Med. Chem. 16, 8187–8195 (2008).

49. F. Liu, Z. Zhang, L. Csanády, D. C. Gadsby, J. Chen, Molecular Structure of the Human CFTR Ion Channel. Cell 169, 85–95.e8 (2017).

50. Z. Zhang, F. Liu, J. Chen, Molecular structure of the ATP-bound, phosphorylated human CFTR. Proc. Natl. Acad. Sci. 115, 12757–12762 (2018).

51. V. Corradi, R.-X. Gu, P. Vergani, D. P. Tieleman, Structure of Transmembrane Helix 8 and Possible Membrane Defects in CFTR. Biophys. J. 114, 1751–1754 (2018).

52. T. C. Hwang, D. N. Sheppard, Molecular pharmacology of the CFTR Cl-channel. Trends Pharmacol. Sci. 20, 448–453 (1999).

53. O. Edelheit, A. Hanukoglu, I. Hanukoglu, Simple and efficient site-directed mutagenesis using two single-primer reactions in parallel to generate mutants for protein structure-function studies. BMC Biotechnol. 9, 61 (2009).

54. A. Goehring, et al., Screening and large-scale expression of membrane proteins in mammalian cells for structural studies. Nat. Protoc. 9, 2574–2585 (2014).

55. Z. Zhang, J. Chen, Atomic Structure of the Cystic Fibrosis Transmembrane Conductance Regulator. Cell 167, 1586–1597.e9 (2016).

56. S. Q. Zheng, et al., MotionCor2: anisotropic correction of beam-induced motion for improved cryo-electron microscopy. Nat. Methods 14, 331–332 (2017).

57. K. Zhang, Gctf: Real-time CTF determination and correction. J. Struct. Biol. 193, 1–12 (2016).

58. S. H. W. Scheres, RELION: Implementation of a Bayesian approach to cryo-EM structure determination. J. Struct. Biol. 180, 519–530 (2012).

59. P. Emsley, K. Cowtan, Coot: model-building tools for molecular graphics. Acta Crystallogr. D Biol. Crystallogr. 60, 2126–2132 (2004).

60. P. D. Adams, et al., PHENIX: a comprehensive Python-based system for macromolecular structure solution. Acta Crystallogr. D Biol. Crystallogr. 66, 213–221 (2010).

61. V. B. Chen, et al., MolProbity: all-atom structure validation for macromolecular crystallography. Acta Crystallogr. D Biol. Crystallogr. 66, 12–21 (2010).

62. B. F. Scharschmidt, E. B. Keeffe, N. M. Blankenship, R. K. Ockner, Validation of a recording spectrophotometric method for measurement of membrane-associated Mg- and NaK- ATPase activity. J. Lab. Clin. Med. 93, 790–799 (1979).

63. M. Dyla, et al., Dynamics of P-type ATPase transport revealed by single-molecule FRET. Nature 551, 346–351 (2017).

64. M. F. Juette, et al., Single-molecule imaging of non-equilibrium molecular ensembles on the millisecond timescale. Nat. Methods 13, 341–344 (2016).

65. F. Qin, Restoration of Single-Channel Currents Using the Segmental k-Means Method Based on Hidden Markov Modeling. Biophys. J. 86, 1488–1501 (2004).

